# Sleep Disruption Selectively Weakens Reactivated Memories

**DOI:** 10.1101/2022.04.10.487316

**Authors:** Nathan W. Whitmore, Ken A. Paller

**Affiliations:** Department of Psychology and Cognitive Neuroscience Program, Northwestern University

## Abstract

A widely accepted view in memory research is that recently stored information can be reactivated during sleep, leading to memory strengthening. Two recent studies have shown this effect can be reversed in participants with highly disrupted sleep. To test whether weakening of reactivated memories can result directly from sleep disruption, in this experiment we varied the intensity of memory reactivation cues, such that some produced sleep arousals. Prior to sleep, participants (local community members) learned the locations of 75 objects, each accompanied by a sound naturally related to that object. Location recall was tested before and after sleep, and a subset of the sounds were presented during sleep. Reactivation with arousal weakened memories, unlike the improvement typically found. We conclude that reactivated memories can be selectively weakened during sleep, and that memory reactivation may strengthen or weaken memories depending on additional factors such as concurrent sleep disruption.

**Statement of Relevance:** The results of this study have implications for both human health and basic psychology. Sleep disorders like apnea are associated with memory problems; our results suggest a possible mechanism where frequent arousal may disrupt the naturally occurring reactivation of memory in sleep. These results also highlight the importance of avoiding brief sleep disruption for good sleep hygiene. Finally, we raise the possibility that reactivation with sleep disruption could be used therapeutically to weaken distressing memories.

Our observation that memories can also be either weakened or strengthened by sleep reactivation has implications for understanding the mechanisms of memory consolidation. In particular, we suggest that sleep memory reactivation may be a reconsolidation-like process with memory restabilization required after reactivation. Our findings also suggest avenues for future experiments; such as using sleep disruption to study the time course of memory reactivation.

## Introduction

A prevalent view in the neuroscience of memory is that reactivation during sleep functions to stabilize memories and reduce forgetting (Born & Wilhelm, 2012; Paller et al., 2020; Pavlides & Winson, 1989; Sirota et al., 2003). This understanding reflects the combination of human research and rodent research, but memory reactivation in human and rodent sleep has been inferred using different methods. In rodent sleep, hippocampal place cells fired in a sequence representing the path the animal took during previous wake (Pavlides & Winson, 1989) This phenomenon has been termed hippocampal replay and has been repeatedly observed (Foster, 2017). In human sleep, EEG activity was shown to reflect the type of information learned before sleep (Schönauer et al., 2017; Schreiner & Staudigl, 2020). Subsequent studies have applied additional methods to yield EEG evidence of memory reactivation during sleep (Belal et al., 2018; Cairney et al., 2018; Wang et al., 2019).

Other experimental approaches have also been used to support the view that memory reactivation during sleep facilitates memory storage, such as with disruptive methods. For example disrupting reactivation in the rodent hippocampus via electrical stimulation impairs learning (Ego-Stengel & Wilson, 2010; Girardeau et al., 2009).In these studies, periods of reactivation were identified by the presence of sharp-wave ripples, a stereotyped discharge in the hippocampus that occurs coincident with place-cell replay (Buzsáki, 2015; Laventure & Benchenane, 2020). Because the same stimulation did not produce memory impairments when delivered outside of sharp-wave-ripple periods, these results suggest that memory was impaired due to disruption of the ripple-associated reactivation process.

In human studies, a procedure known as targeted memory reactivation (TMR) has been used to probe the effects of memory reactivation during sleep (Oudiette & Paller, 2013). In a typical TMR study, to-be-learned information is associated with a sensory stimulus (e.g., a specific sound), which is subsequently presented during sleep without waking the participant. A recent meta-analysis has substantiated these sorts of results (Hu et al., 2020). In within-subjects designs, memory is measured both before and after a sleep period, and some items (but not others) are reactivated during sleep. Reactivated items are typically remembered better on the post-sleep test than items not reactivated. In between-subject studies, participants who receive reactivation during sleep perform better on post-sleep tests than subjects who received irrelevant sounds during sleep.

The TMR procedure relies on the premise that sensory stimulation can be delivered during sleep without producing awakening or arousal from sleep. However, sleep may indeed be disrupted under some circumstances. Göldi and Rasch (2019) described a TMR procedure in participants’ homes, unsupervised by laboratory personnel, and they found that when participants reported that reactivation cues disturbed their sleep, reactivated items were remembered less well than items not reactivated. However, there were no measures of sleep physiology to assess the sleep disruption. Subsequently, in a laboratory study, Whitmore and colleagues (2022) found that participants with shallow sleep and large numbers of arousals evident in EEG recordings did not show the normal benefits of TMR, and in some cases showed weakening of reactivated memories.

What might explain these effects? We hypothesized that reactivation combined with sleep disruption introduces errors into memory traces. This model rests on two claims. The first claim is that memory engrams can be modified when memories are reactivated during sleep, thereby allowing for strengthening and consolidation. The second claim is that if sleep does not remain undisturbed during reactivation, but is instead disrupted, consolidation can go awry, inducing errors in the reactivated memory trace.

An initial prediction of this hypothesis is that memory reactivation accompanied by sleep disruption should produce forgetting, unlike reactivating a memory without sleep disruption. To test this prediction, we developed an experiment in which we reactivated memories during sleep with a varying amount of sleep disruption.

### Open Practices Statement

The study reported in this article was not preregistered. Data and code are available from the corresponding author upon reasonable request.

## Methods

Participants (*N*=24) were a convenience sample of 11 males and 13 females 18-30 years old (mean=21.6, SD=3.79) recruited from the Northwestern University population and surrounding community. Inclusion criteria included being between 18-35 years old, self-reporting being able to sleep in the afternoon in the lab, and currently not having a sleep disorder. Our target sample size was selected to be comparable to that used in other sleep reactivation experiments in and outside our lab. To increase the likelihood of falling asleep, participants were asked to get 1 hour less sleep than normal the night before and to avoid nicotine and caffeine on the day of the study. After participants arrived in the lab and gave written informed consent, the following six phases transpired: initial learning, bioelectric recording setup, pre-sleep memory test, sleep, post-sleep memory test, and a test to assess which sounds (if any) were heard during the nap. Figure 1 shows an overview of the procedure, which was approved by the Northwestern University Institutional Review Board.

**Figure 1:**
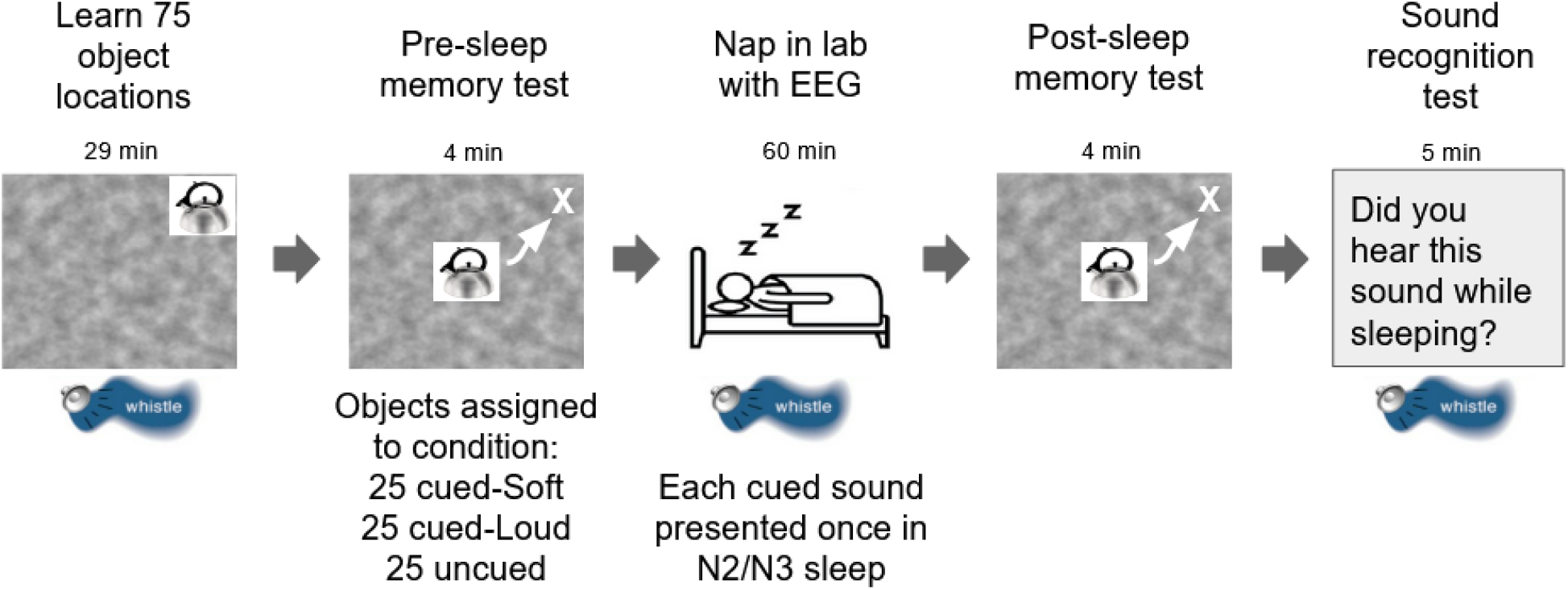
Illustration of experimental procedure, with mean duration for each phase. Participants first learned locations of 75 objects paired with sounds on a Perlin-noise background. Bioelectric recording set-up was next (not shown, mean time 29 min). Participants then took a memory test where they moved objects (illustrated here by white arrow and X) from the center to the correct location. Following memory testing, participants slept in the lab while object sounds were presented in N2 and N3 sleep. Following sleep, participants performed a second memory test, identical to the first. Finally, we played each of the 75 sounds and participants indicated whether or not they had heard each sound during sleep.

### Procedure

Participants learned the locations of 75 pictures of common objects presented individually on a Perlin noise background (Figure 1). The learning consisted of two parts. In the first part, participants were shown each object in its correct location and then immediately asked to move the object from the center of the screen to the correct location. Participants were shown the correct location after placement and received visual feedback (either “correct” or “incorrect” with the correct location of the object shown). Correct responses were those placed less than 3 cm from the correct location. The sound associated with each object was played when it first appeared on the screen and when participants made a correct response.

After performing this procedure for all 75 objects, participants began the second part, which required learning to criterion. Objects were presented in the center of the screen and participants were asked to move them to the correct location. As in the first part, sounds were presented when the object first appeared and when the participant made a correct response. Objects were presented in a random order, constrained so that the same object could not be shown twice in a row unless it was the only object remaining. After placing the object, participants received feedback in the same manner as during the first part. If the participant placed the object center within 3 cm of the correct location, the criterion was considered achieved and the object was not shown again; otherwise, the object was included in the rotation. Learning ended when the participant correctly placed all the objects.

### Bioelectric recording

Following learning, we attached bioelectric recording electrodes for EEG, EMG, and EOG (electroencephalography, electromyography, and electro-oculography). Data were recorded using a Neuroscan Synamps2 system with 26 scalp channels plus horizontal and vertical EOG, and chin EMG. Data were recorded at 1000 Hz with a high-pass filter at 0.1 Hz and a low-pass filter at 100 Hz.

### Pre-sleep test

After bioelectric recording setup, participants completed a pre-sleep memory test. In this test, objects were presented in random order and the participant attempted to place each object at its correct location. No feedback or sounds were presented during the pre-sleep test and each object was tested once.

After the test was complete, objects were divided into three sets comprising 25 to be cued with loud sounds during sleep, 25 to be cued with soft sounds, and 25 not cued. Objects were assigned to sets so as to match pre-sleep memory performance across sets. In this procedure, the objects were first ranked by accuracy and sequentially assigned to sets, so that each set received an equal mix of high-, medium-, and low-accuracy objects.

### Sleep period

Participants slept on a futon in the same chamber where they completed the behavioral tasks. When participants reached stage N2 sleep (determined by the experimenter’s real-time sleep staging), their initial arousal threshold was determined by presenting a probe sound (bike bell) not related to the memory task. If the sound did not elicit an arousal, the intensity was raised and the sound presented again, repeating this procedure until an arousal occurred. The intensity that prompted an arousal was used as the initial intensity for the sounds in the loud set.

After finding the arousal threshold, we waited for the participant to return to stable N2 sleep and then began presenting the cue sounds. Sounds were presented in random order, with loud and soft sounds intermixed. Loud sounds were presented at approximately 43 dBa; with intensity continually adjusted to reliably produce brief arousals but avoid prolonged awakenings, defined as more than a minute of wake or N1 following the cue. Quiet sounds were presented at low intensity (approx. 28 dBa) and adjusted to avoid arousal. The mean intensity of quiet sounds was 33% of the initial arousal threshold [SEM=4.3%] and 7% of the mean intensity of loud sounds [SEM=0.6%]. Decibel values were determined by testing using Decibel X on a Redmi Note 9 placed at the location of the participant’s head.

Sounds were presented with at least a 10-s interstimulus interval. If a sound triggered an arousal, cueing was paused until the participant returned to stable N2 or N3 sleep. Each sound was presented only once, allowing us to correlate each object’s spatial memory fate with the sleep physiology surrounding sound presentation. After all sounds were presented, the participant was allowed to sleep for 5 min before they were awakened.

Immediately after the participant awoke, we informed them that we had played sounds during the nap and asked them if they remembered hearing any sounds. This was the first time that participants were explicitly told that sounds would be presented during the nap.

### Post-sleep test

Participants performed the post-sleep memory test, which was identical to the pre-sleep test, approximately 5 min after awakening.

### Sound recognition test

Following the post-sleep memory test, we presented the 75 object-associated sounds one at a time. Participants were asked to indicate whether they heard each sound during their sleep, with three possible responses: Definitely yes, possibly, and definitely no.

### Sound intensity analysis

We defined the forgetting ratio (FR) across the sleep period as post-sleep error (in cm) / pre-sleep error (in cm). We computed a 3-level repeated-measures ANOVA to compare FR for cued-loud objects, cued-soft objects, and uncued objects. We also computed a two-tailed repeated-measures *t* test to compare FR for all cued objects to uncued objects.

### EEG data processing

EEG data were analyzed in EEGLAB 2020.0 (Delorme & Makeig, 2004). Prior to analysis, we visually inspected the data and replaced EEG channels with poor signal quality using interpolation. No other data pre-processing or cleaning was performed. Data from one participant were excluded from arousal and EEG spectrum analyses due to a technical failure that prevented stimulus times from being recorded.

### Arousal analysis

To measure the effects of EEG arousal on memory, we performed offline manual scoring to classify each sound cue as arousal-provoking or non-arousal-provoking. During this classification, the rater was blinded to the type of cue (loud versus soft). The rater examined a segment of time from 10 s before to 10 s after each cue, and scored the cue as arousal-provoking if an arousal meeting the AASM criteria (Iber et al., 2007) occurred in the 10 s after the cue. We computed FR for three conditions: cued objects that produced an arousal, cued objects that did not produce an arousal, and uncued objects. We then tested whether FR differed for objects cued with arousal, objects cued with no arousal, and uncued objects in a 3-level repeated-measures ANOVA.

### Effects of cue perception on memory fate

To assess the impact of consciously perceiving sounds during sleep on memory fate, we compared FR for objects where participants reported hearing the associated sound to FR for objects where participants did not report hearing the sound. In this analysis, sounds were considered heard if the participant reported definitely or possibly hearing them during the sleep period. We tested whether FR differed for cued-heard and cued-unheard objects using a two-tailed repeated-measures *t* test.

### EEG predictors of memory fate

To identify EEG features associated with enhancement/weakening of memory, we correlated FR for each cued object with the power spectrum from the 10 s before the cue. Spectra were computed from electrode Cz using the spectopo function in EEGLAB v2020.0. Each spectrum consisted of 21 2.5-Hz-wide frequency bins, spanning 0-52.5 Hz. Correlations were performed using a mixed model in JMP 15, with participant as a random factor and spectral power in each bin as a fixed factor.

## Results

### Spatial recall accuracy was matched across conditions before sleep, and declined after sleep, especially for cued objects

At the pre-sleep test, participants recalled locations with a mean error of 6.8 cm [SEM=0.52]. Recall accuracy did not differ between cued-loud, cued-soft, and uncued objects [F(2,46)=1.22, *p*=0.3].

Recall error increased at the post-sleep test to a mean of 7.47 cm [SEM=0.53], which was significantly worse than pre-sleep [*t*(23)=2.89, *p*=0.008]. As shown in Figure 2, recall was less accurate after sleep in all three conditions and there was a larger increase in error (FR) for all cued objects combined compared to uncued objects [*t*(23) = 2.39, *p* = 0.025]. While this effect was significant in our planned comparison of all cued objects versus uncued objects, no significant differences were found in the omnibus ANOVA comparing cued-loud, cued-soft, and uncued sets [F(2,46)=1.64, *p*=0.21].

**Figure 2:**
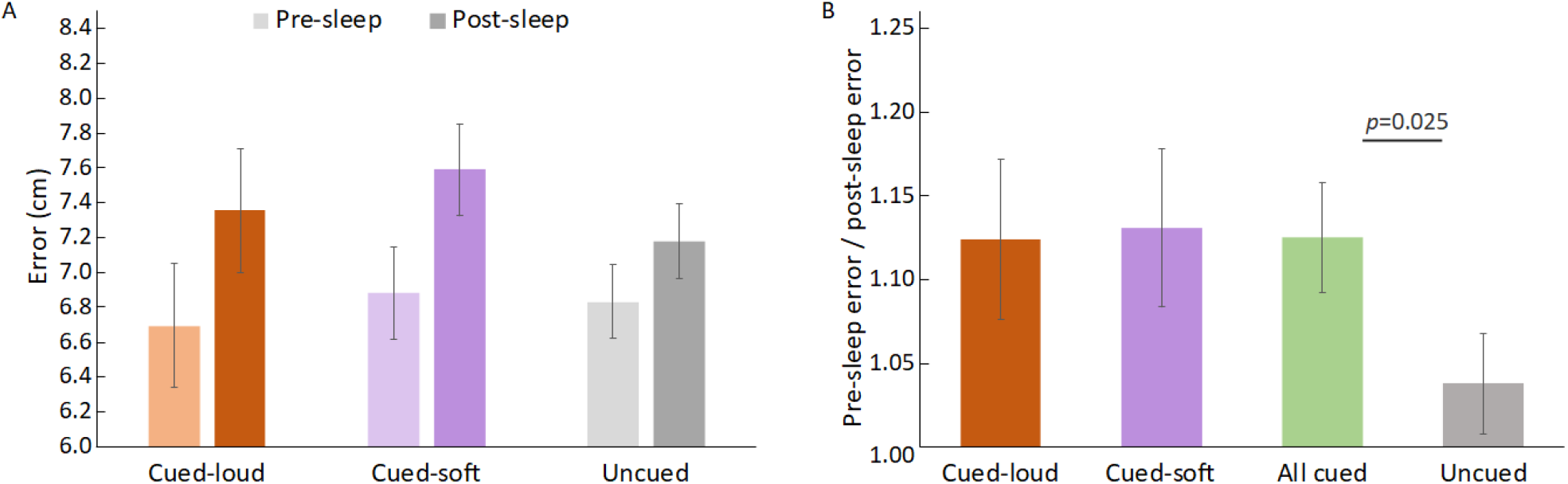
(A) Mean spatial error in centimeters for all conditions on pre-and post-sleep tests. Error bars represent the SEM for the post-sleep error minus pre-sleep error within subjects. (B) Forgetting Ratio (fold change in spatial error) for all conditions.

### Forgetting was increased when cues generated an arousal

To understand the effects of arousal during sleep on reactivation, we categorized cued objects according to whether arousal was apparent in post-stimulus EEG recordings. Arousals were common for both types of sound. Arousals occurred following 43% of the soft sounds and 59% of the loud sounds.

As shown in Figure 3, forgetting was greater for objects that were cued with arousal than for uncued objects. On the pre-sleep test, recall did not differ by condition [F(2,44)=2.05, *p*=0.14] whereas differences were apparent after sleep. An ANOVA comparing FR for uncued objects, objects cued with arousal, and objects cued without arousal revealed a significant effect [F(2,44)=3.51, *p*=0.04]. Post-hoc testing using two-tailed t tests of all pairs with Bonferroni correction (alpha per test=0.017) showed a significant difference between uncued objects cued with arousal and uncued objects [*t*(22)=3.28, *p*=0.003].

**Figure 3:**
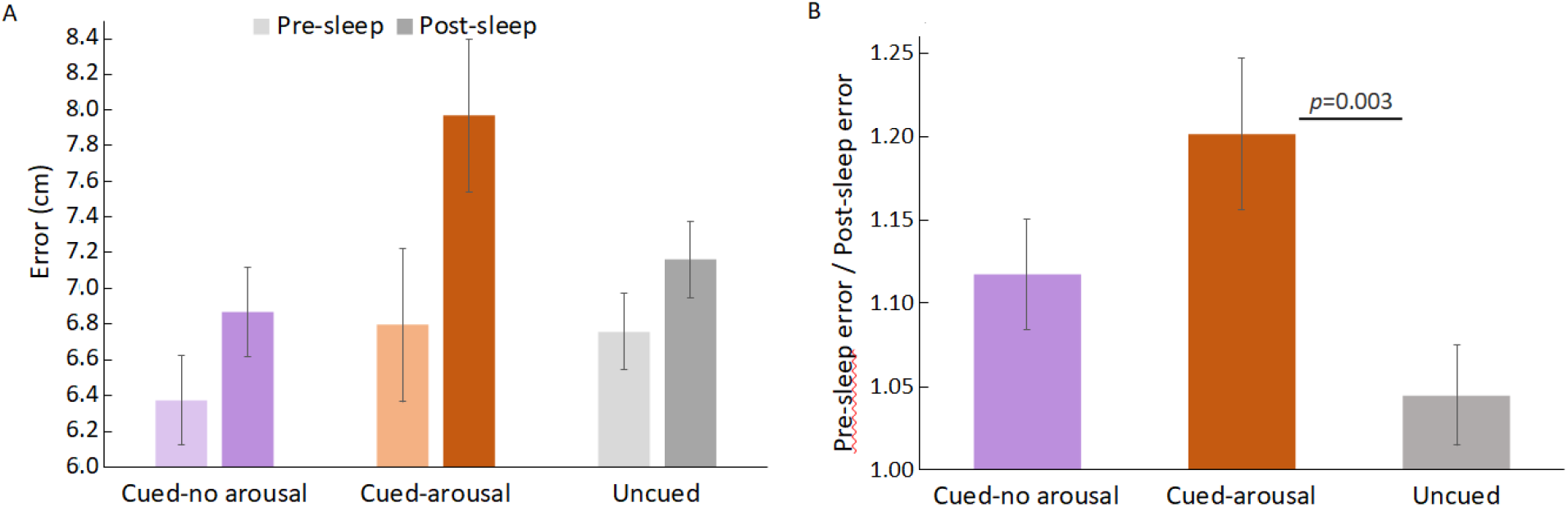
(A) Mean error for uncued objects and objects cued with/without arousal. Error bars represent the SEM for post-sleep error minus pre-sleep error within subjects. (B) Forgetting ratio for these three conditions.

### Pre-cue alpha power predicts the effect of cueing on memory

To identify EEG features that predicted the effects of cueing, we tested whether EEG spectral power at Cz in the 10 s prior to a cue predicted the memory fate of the associated object. We divided the spectrum into 21 2.5-Hz wide bins, spanning 0-52.5 Hz. Increased power in the high alpha bin (10-12.5 Hz) before the cue was associated with more forgetting of the cued object [*t* (642.5)=2.37, *p*=0.018]. However, following FDR multiple-comparisons correction this correlation was nonsignificant (*p*=0.38).

### Subjective recall of sleep cues did not influence memory performance

Because participants sometimes report hearing cue sounds during sleep, we examined possible relationships between subjective reports of hearing a sound during sleep and changes in the corresponding object-location memories. At the end of the experiment, participants were presented with each cue sound and asked if they remembered hearing it while sleeping. On average, participants reported definitely hearing 11% of the sounds played during sleep [SEM = 3%] and possibly hearing 28% of the sounds [SEM = 4%]. We combined these two categories for this analysis. There was no significant difference in FR [*t*(23)=0.88,*p*=0.39] between cued objects with sounds heard during sleep [mean FR=1.25, SEM=0.17], and cued objects with sounds not heard during sleep [mean FR=1.09, SEM=0.04].

## Discussion

In this experiment, we observed that reactivating memories in the context of disrupted sleep produced a selective forgetting effect, where spatial recall accuracy after sleep was decreased for the reactivated objects. These results support our hypothesis that sleep disruption can reverse the typical accuracy-enhancing effect of TMR.

Surprisingly, we did not observe a difference in memory outcomes for objects cued with soft sounds and objects cued with loud sounds. One potential explanation is that many soft sounds also triggered an arousal; the presence or absence of an arousal may be more relevant to memory than the absolute sound volume. Supporting this hypothesis, objects cued with arousal had a significantly higher forgetting ratio than uncued objects and slightly (but not significantly) higher forgetting ratio than objects cued without arousal. Soft sounds may be more likely to trigger arousals when sleep is repeatedly interrupted, as it was in this experiment.

Another surprising observation was that we did not observe a memory-enhancing effect of TMR; objects reactivated without arousal were comparable to uncued objects, with a slight but not significant increase in error. The lack of benefit for objects without arousal might indicate that the effects of sleep disruption can “spread” and affect not only memories reactivated with arousal but also related memories or memories reactivated around the same time.

Our results also suggest that sleep state prior to a cue can influence the fate of a memory. When cues were presented during periods of sleep with relatively high alpha power, the corresponding object locations were recalled less accurately. Given that alpha power during sleep is thought to reflect sleep depth and arousability (McKinney et al., 2011), a reasonable interpretation is that cues presented in periods of light sleep are more likely to trigger arousal and weakening.

Finally, we found that participant reports of hearing cues did not significantly predict memory fate. This result contrasts with those of Göldi and Rasch (2019), where only participants who reported sleep disturbed by cues showed worsening of memory induced by TMR. This difference across experiments may reflect differences in the questions used to assess sleep. Göldi and Rasch asked participants whether the sounds woke them, whereas we asked if participants remembered hearing each sound. Asking about waking (as opposed to memory for hearing the sound) may therefore be a better proxy of TMR-induced arousal.

Our results parallel the effects of disrupting memory reconsolidation. While retrieval during wake typically strengthens memory (Roediger & Butler, 2011), Misanin and colleagues (1968) showed that a retrieval cue followed by electroconvulsive stimulation produced forgetting of the reactivated information in rats. Similarly, retrieval while protein synthesis is inhibited produces forgetting (Nader et al., 2000). Because both of these interventions produce amnesia by disrupting early consolidation (Haubrich & Nader, 2018), these results imply that consolidation processes are required after retrieval of a memory to prevent forgetting. If consolidation after retrieval is disrupted, the memory becomes weakened. Paralleling this idea, TMR with uninterrupted sleep may allow a full consolidation process to occur, thereby strengthening memory, whereas interrupting sleep may prevent stabilization from occurring. Cueing followed by sleep disruption may thus produce a destabilizing effect, leading to memories that tend to be less accurate upon awakening.

An open question is to what extent these effects may occur in other settings. Our study examined sleep during a nap of limited duration using cues to reactivate specific memories. Future studies could explore whether these same effects occur in overnight sleep, or with spontaneous memory reactivation, as well as how sleep disruption associated with sleep disorders may affect memory.

Our results add to the literature on memory processing during sleep by showing that it is possible for memory reactivation to lead not only to strengthening, but also to weakening. Weakening may be particularly perpetuated when sleep is disrupted. This finding raises several interesting possibilities. The ability to produce both strengthening and weakening may prove useful in understanding the discrete processes that operate during memory reactivation in sleep. The results also suggest potential practical applications, such as using TMR to weaken memories of traumatic events, which could be explored in future research.

## Author Contributions

N.W.W. and K.A.P. designed the experiment. N.W.W. collected, analyzed, and interpreted data. K.A.P. supervised the project. N.W.W. and K.A.P. wrote the manuscript.

## Acknowledgements

We are grateful for funding from NIH (R01-NS112942, T32-NS047987, and T32-MH06756) and a McKnight Memory and Cognitive Disorders Award from the McKnight Endowment Fund for Neuroscience.

## References

Belal, S., Cousins, J., El-Deredy, W., Parkes, L., Schneider, J., Tsujimura, H., Zoumpoulaki, A., Perapoch, M., Santamaria, L., & Lewis, P. (2018). Identification of memory reactivation during sleep by EEG classification. NeuroImage, 176, 203–214. https://doi.org/10.1016/j.neuroimage.2018.04.029

Born, J., & Wilhelm, I. (2012). System consolidation of memory during sleep. Psychological Research, 76(2), 192–203. https://doi.org/10.1007/s00426-011-0335-6

Buzsáki, G. (2015). Hippocampal sharp wave-ripple: A cognitive biomarker for episodic memory and planning. Hippocampus, 25(10), 1073–1188. https://doi.org/10.1002/hipo.22488

Cairney, S. A., Guttesen, A. á V., El Marj, N., & Staresina, B. P. (2018). Memory consolidation is linked to spindle-mediated information processing during sleep. Current Biology, 28(6), 948–954.e4. https://doi.org/10.1016/j.cub.2018.01.087

Delorme, A., & Makeig, S. (2004). EEGLAB: An open source toolbox for analysis of single-trial EEG dynamics including independent component analysis. Journal of Neuroscience Methods, 134(1), 9–21. https://doi.org/10.1016/j.jneumeth.2003.10.009

Ego-Stengel, V., & Wilson, M. A. (2010). Disruption of ripple-associated hippocampal activity during rest impairs spatial learning in the rat. Hippocampus, 20(1), 1–10. https://doi.org/10.1002/hipo.20707

Foster, D. J. (2017). Replay comes of age. Annual Review of Neuroscience, 40(1), 581–602. https://doi.org/10.1146/annurev-neuro-072116-031538

Girardeau, G., Benchenane, K., Wiener, S. I., Buzsáki, G., & Zugaro, M. B. (2009). Selective suppression of hippocampal ripples impairs spatial memory. Nature Neuroscience, 12(10), 1222–1223. https://doi.org/10.1038/nn.2384

Göldi, M., & Rasch, B. (2019). Effects of targeted memory reactivation during sleep at home depend on sleep disturbances and habituation. NPJ Science of Learning, 4, 5. https://doi.org/10.1038/s41539-019-0044-2

Haubrich, J., & Nader, K. (2018). Memory reconsolidation. In R. E. Clark & S. J. Martin (Eds.), Behavioral Neuroscience of Learning and Memory (pp. 151–176). Springer International Publishing. https://doi.org/10.1007/7854_2016_463

Hu, X., Cheng, L. Y., Chiu, M. H., & Paller, K. A. (2020). Promoting memory consolidation during sleep: A meta-analysis of targeted memory reactivation. Psychological Bulletin, 146(3), 218–244. https://doi.org/10.1037/bul0000223

Iber, C, Ancoli-Israel, S, Chesson, A, & Quan, SF. (2007). The AASM manual for the scoring of sleep and associated events: Rules, terminology, and technical specifications (1st ed.). American Academy of Sleep Medicine.

Laventure, S., & Benchenane, K. (2020). Validating the theoretical bases of sleep reactivation during sharp-wave ripples and their association with emotional valence. Hippocampus, 30(1), 19–27. https://doi.org/10.1002/hipo.23143

McKinney, S. M., Dang-Vu, T. T., Buxton, O. M., Solet, J. M., & Ellenbogen, J. M. (2011). Covert waking brain activity reveals instantaneous sleep depth. PLOS ONE, 6(3), e17351. https://doi.org/10.1371/journal.pone.0017351

Nader, K., Schafe, G. E., & Le Doux, J. E. (2000). Fear memories require protein synthesis in the amygdala for reconsolidation after retrieval. Nature, 406(6797), 722–726. https://doi.org/10.1038/35021052

Oudiette, D., & Paller, K. A. (2013). Upgrading the sleeping brain with targeted memory reactivation. Trends in Cognitive Sciences, 17(3), 142–149. https://doi.org/10.1016/j.tics.2013.01.006

Paller, K. A., Mayes, A. R., Antony, J. W., & Norman, K. A. (2020). Replay-based consolidation governs enduring memory storage. The Cognitive Neurosciences, Sixth Edition, MIT Press. https://par.nsf.gov/biblio/10187208-replay-based-consolidation-governs-enduring-memory-storage

Pavlides, C., & Winson, J. (1989). Influences of hippocampal place cell firing in the awake state on the activity of these cells during subsequent sleep episodes. The Journal of Neuroscience: The Official Journal of the Society for Neuroscience, 9(8), 2907–2918.

Roediger, H. L., & Butler, A. C. (2011). The critical role of retrieval practice in long-term retention. Trends in Cognitive Sciences, 15(1), 20–27. https://doi.org/10.1016/j.tics.2010.09.003

Schönauer, M., Alizadeh, S., Jamalabadi, H., Abraham, A., Pawlizki, A., & Gais, S. (2017). Decoding material-specific memory reprocessing during sleep in humans. Nature Communications, 8(1), 15404. https://doi.org/10.1038/ncomms15404

Schreiner, T., & Staudigl, T. (2020). Electrophysiological signatures of memory reactivation in humans. Philosophical Transactions of the Royal Society B: Biological Sciences, 375(1799), 20190293. https://doi.org/10.1098/rstb.2019.0293

Sirota, A., Csicsvari, J., Buhl, D., & Buzsáki, G. (2003). Communication between neocortex and hippocampus during sleep in rodents. Proceedings of the National Academy of Sciences, 100(4), 2065–2069. https://doi.org/10.1073/pnas.0437938100

Wang, B., Antony, J. W., Lurie, S., Brooks, P. P., Paller, K. A., & Norman, K. A. (2019). Targeted memory reactivation during sleep elicits neural signals related to learning content. Journal of Neuroscience, 39(34), 6728–6736. https://doi.org/10.1523/JNEUROSCI.2798-18.2019

Whitmore, N. W., Bassard, A. M., & Paller, K. A. (2022). Targeted memory reactivation of face-name learning depends on ample and undisturbed slow-wave sleep. Npj Science of Learning, 7(1), 1–6. https://doi.org/10.1038/s41539-021-00119-2

